# The effect of salt on the dynamics of CoV-2 RBD at ACE2

**DOI:** 10.1101/2020.10.09.333948

**Authors:** E. K. Peter, A. Schug

## Abstract

In this article, we investigate the effect of electrolytes on the stability of the complex between the coronavirus type 2 spike protein receptor domain (CoV-2 RBD) and ACE2, which plays an important role in the activation cascade at the viral entry of CoV-2 into human cells. At the cellular surface, electrolytes play an important role, especially in the interaction of proteins near the membrane surface. Additionally, the binding interface of the CoV-2 RBD - ACE2 complex is highly hydrophilic. We simulated the CoV-2 RBD - ACE2 complex at varying salt concentrations over the concentration range from 0.03 M to 0.3 M of calcium and sodium chloride over an individual simulation length of 750 ns in 9 independent simulations (6.75 *µs* total). We observe that the CoV-2 RBD - ACE2 complex is stabilized independent of the salt concentration. We identify a strong negative electrostatic potential at the N-terminal part of CoV-2 RBD and we find that CoV-2 RBD binds even stronger at higher salt concentrations. We observe that the dynamics of the N-terminal part of CoV-2 RBD stabilize the protein complex leading to strong collective motions and a stable interface between CoV-2 RBD and ACE2. We state that the sequence of CoV-2 RBD might be optimized for a strong binding to ACE2 at varying salt concentrations at the cellular surface, which acts as a key component in the activation of CoV-2 for its viral entry.

**SIGNIFICANCE:** A novel coronavirus, coronavirus type 2 (CoV-2), was identified as primary cause for a worldwide pandemic of the severe acute respiratory syndrome (SARS CoV-2). The CoV-2 spike protein is a major target for the development of a vaccine and potential strategies to inhibit the viral entry into human cells. At the cellular surface, CoV-2 activation involves the direct interaction between ACE2 and CoV-2 RBD. At the cellular surface, electrolytes play an important role, especially in the interaction of proteins near the membrane surface. We thus investigate the effect of ion conditions on the interaction of the CoV-2 RBD - ACE2 complex and find stabilizing effects. We speculate that CoV-2 RBD is optimized for strong binding to ACE2 at varying salt concentrations.

## INTRODUCTION

A novel class of coronaviruses appearing in November 2019 and 2020 (CoV-19 and CoV-2) are the major cause severe acute respiratory syndrome (SARS). The global SARS pandemic reached an unprecedented rising number of 28.637.952 infections and 917.417 deaths worldwide (1). Till the onset of the last SARS-CoV pandemic in 2003 (2, 3), the potential of transfection of SARS-CoV between different organisms was strongly underestimated. Before SARS-CoV 2003, CoV was considered as a class of viruses that cause mild respiratory diseases, such as cold or infectious bronchitis (4). Except other cases, e.g. Bovine CoV (BCoV), which can be transfected from cattle to turkeys and humans, SARS-CoV possesses a transfectious potential between cats, racoons, humans, macaques and civet cats (5). Mainly due to political reasons, a clear localization of the first cluster of infections and the transfection from animals to humans that triggered the global pandemic still remains speculative. However, it is widely accepted that the first cluster of infections occurred at the Huanan sea food marked due to animal-to-human transfection in Wuhan, China in 2019 (6–8). CoV belong to the class of *Coronavirinae*, which contain a characteristic pattern of spike (S) proteins surrounding the viral capsid. The CoV S proteins act as activating entities for the viral entry into target cells and remain the major target for a potential vaccination strategy against SARS CoV-2. The S-protein forms trimers at the protrusions of the virus and comprises two functional subunits : S1 and S2. In the sequence of events along the viral entry, the S1 unit of the spike (S) protein facilitates the attachment of the virus at the surface of the target cell (9). The S2 subunit, responsible for membrane fusion, employs TMPRSS2 for S protein priming, while it uses ACE2 as entry receptor for membrane fusion (10–15). Notably, one of the key factors for its infectious potential for humans is the high conservation of ACE2 in different mammalian organisms (16), which allows its transmission from animals to humans. The receptor binding domain (RBD) of the S1 subunit contains five antiparallel beta strands, while alpha-helical and loop motifs form the connecting entities between the beta sheets. Between two of the central beta sheet motifs, an extended insertion forms the receptor binding motif (RBM), which binds to ACE2 at its N-terminal helix (11, 17–20). Among a large number of potential targets, the inhibition of the direct interaction between ACE2 and the S-protein (SARS CoV-S) provides a suitable strategy to prevent the membrane fusion of CoV-2 and the viral entry into human cells (21, 22). The structure of CoV-2 RBD in complex with ACE2 has been characterized in an X-ray structure, which provides insight into the viral activation process at the target cell (17). Surprisingly, the interface between CoV-2 RBD and ACE2 is comprised of hydrophilic interactions, while a hydrophobic patch is located at the turn region of CoV-2 RBD, which lies on the opposite side of the CoV-2 RBD - ACE2 interface.

To gain access into dynamics properties of biomolecular properties, molecular dynamics simulations have been a valuable addition to experimental investigations. Examples include coarse-grained simulations of protein folding (23–25), conformational transitions(26), RNA(27, 28), complementing experimental information(29–32), or simulations of huge systems (33, 34).

In a enhanced correlation guided MD (CORE-MD) (35) simulation of the assembly process of CoV-2 RBD and ACE2, we observed the formation of a hydrophobic interface, where the hydrophilic region of CoV-2 RBD was rotated away from ACE2 due to electrostatic forces (36). We attributed two different effects to that observation : (1) The effect of an implicit solvent environment in the CORE-MD simulation and (2) a strong electrostatic repulsion between the two domains. We concluded that surrounding electric fields due to salts present in the CoV-RBD - ACE2 system might play a major role in the interaction between CoV-2 RBD and ACE2.

In this article, we present our results from simulations of the CoV-2 RBD - ACE2 system at different ionic strengths. We performed simulations of the CoV-2 RBD - ACE2 complex at varying salt concentrations over the concentration range from 0.03 M to 0.3 M of calcium and sodium chloride over an individual simulation length of 750 ns in 9 independent simulations (6.75 *µs* total). We find that the CoV-2 RBD - ACE2 complex is stabilized independent of the salt concentration. We observe a strong negative electrostatic potential at the N-terminal part of CoV-2 RBD and we see that CoV-2 RBD binds even stronger at higher salt concentrations. We find that the dynamics of the N-terminal part of CoV-2 RBD stabilize the protein complex leading to strong collective motions and a stable interface between CoV-2 RBD and ACE2. We state that the sequence of CoV-2 RBD might be optimized for a strong binding to ACE2 at varying salt concentrations at the cellular surface, which acts as a key component in the activation of CoV-2 for its viral entry.

## METHODS

### Simulation parameters and system setup

We used the GROMACS simulation package version 4.6. for the simulations and the trajectory analysis (37). For all simulations, we used the X-ray structure of the CoV-2 RBD - ACE2 complex (PDB: 6M0J (17)) and centered the system in a box with dimensions 8.076 × 9.108 × 13.543 *nm*^3^ (see Figure 1). We filled each system with 29043 TIP4P waters. Using the program *gmx genion*, we replaced the water molecules with ions, which were inserted randomly across the system. We used the AMBER99SB forcefield to describe the interactions in the system (38). To account for polarizability effects in the system described by non-polarizable point charges, we applied the charge scaling procedure by a factor equal 0.7 for all ionic charges (39). We simulated the systems with sodium chloride and calcium chloride within the concentration range from 0.03 (*NaCl*), 0.06 (*CaCl*_2_) to 0.3 M. We used particle mesh Ewald (PME) electrostatics with a real space cutoff of 0.8 nm and calculated the van der Waals interactions with a shift function using the same cutoff. We used a timestep equal 0.001 ps and applied stochastic velocity rescaling combined with the berendsen barostat to simulate in the NPT-ensemble at a temperature equal 300 K and a pressure equal 1 bar (40). The list of simulations in given in Table 1.

**Table 1:** MD and CORE-MD enhanced sampling simulations which were performed in this study to investigate salt effects on the stability of the CoV-2-RBD - ACE2 complex.

| Simulation | Length | System |
| --- | --- | --- |
| 5 independent simulations, NPT-MD | 750 ns | $CaCl_2$ $c = 0.03 - 0.3$ M |
| 4 independent simulations, NPT-MD | 750 ns | $NaCl$ $c = 0.06 - 0.3$ M |

**Figure 1:**
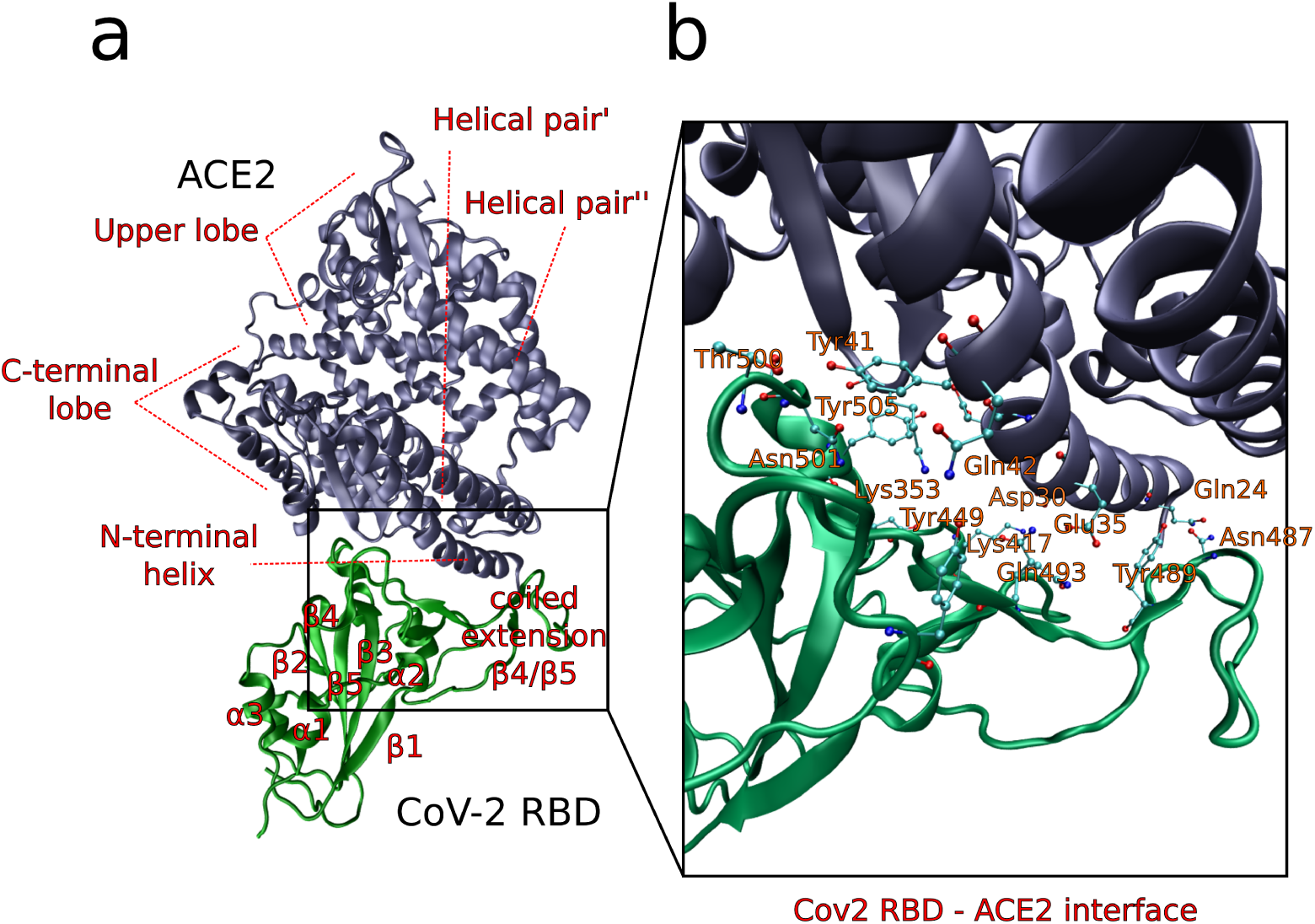
Experimental structure of the CoV-2 RBD - ACE2 complex (PDB: 6m0j (17)). (a) Complete view of the CoV-2 RBD - ACE2 complex. (b) Hydrophilic interface between CoV-2 RBD - ACE2 with residue names depicted with orange letters.

We determined the relative log likelihood of the pair interactions between CoV-2 RBD and ACE2 using (41, 42) :

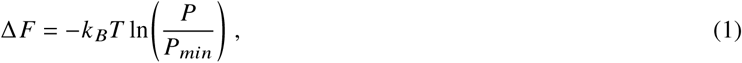

where *P* stands for the probability and *P*_*min*_ is the minimum reference value. We used in-house programs for the calculation of the distance dependent probabilities.

## RESULTS AND DISCUSSION

### Properties of the CoV-2 RBD - ACE2 complex at different salt concentrations

For the CoV-2 RBD - ACE2 complex at different concentrations of *CaCl*_2_, we observe that the RMSD to the crystal structure (PDB : 6m0j) varies between 0.2 to 0.5 nm, while we only observe a stronger fluctuation for the simulation at 0.08 M *CaCl*_2_ at a time range from 600 to 750 ns (see Figure 2 a). In average, the fluctuations of the same complex are larger for the same complex immersed in *NaCl*. We find that the average RMSD is about 0.1 nm higher than for the case with *CaCl*_2_. In general, we observe that the RMSD is at 0.55 at low salt concentrations, while the lowest RMSD is reached at a concentration of 0.3 M (see Figure 2 b). We then analyzed the averaged binding patterns between CoV-2 RBD and ACE2 as function of the index of ACE2 and CoV-2 RBD (see Figure 2 c-f). All simulations show that there is no perturbation of the binding between both proteins independent from the salt concentration. We find that the contacts between the N-terminal helix of ACE2 and the coiled extension *β*4 / *β*5 (ce *β*4 / *β*5) are conserved (see Figure 2 c, d). We observe an analogous behavior for the contacts between residues of ACE2 with indices ranging from #300 to #380 and CoV-2 RBD. We observe stepwise variations in the residue range between # 100 and #290 as well as in the residue range between #400 and #520, which we interpret as different fluctuation ranges due to the salt present in the system. We continued to analyze the contact patterns of the complex as function of the residue index of CoV-2 RBD (see Figure 2 e, f). In these graphs, we observe that the N-terminal region between *α*1 and *β*4 between residues # 616 and # 700 shows a salt concentration dependent behavior, with a rise in the distance by approximately 0.3 nm in relation to ACE2. We find an analogous behaviour at the C-terminus of CoV-2 RBD, where the distance range increases by 0.5 nm, while the binding of the coiled extension to ACE2 remains approximately stable.

**Figure 2:**
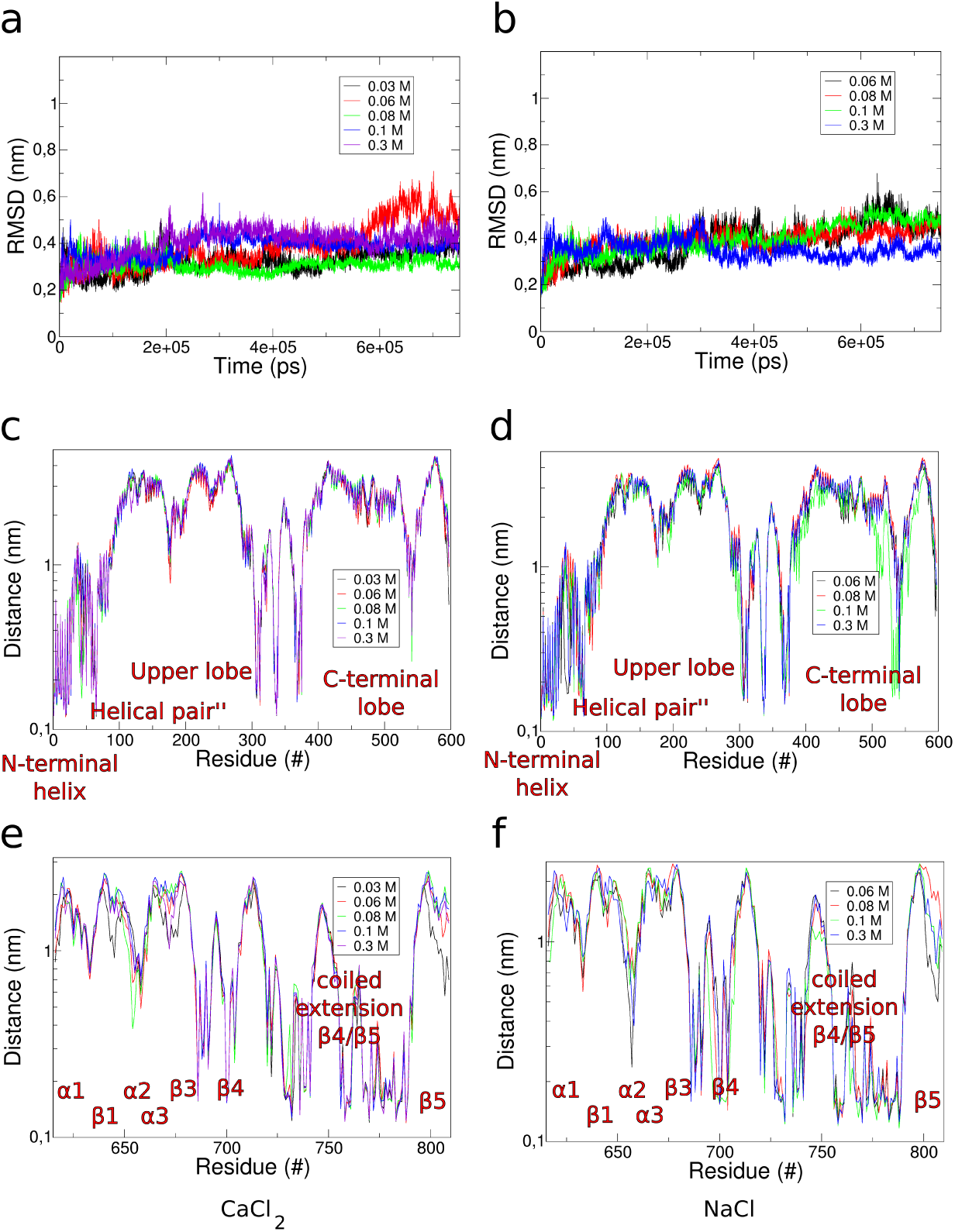
(a) *RMSD*_*C a*− *C a*_ to the crystal structure (PDB : 6m0j) as function of simulation time at different salt concentrations of *CaCl*_2_ ranging from 0.03 to 0.3 M. (b) *RMSD*_*C a C a*_ to the crystal structure (PDB : 6m0j) as function of simulation time at different salt concentrations of *NaCl* ranging from 0.06 to 0.3 M. (c) Minimum distances between ACE2 and CoV-2 RBD as function of the residue index of ACE2 averaged over the trajectories from simulations at different concentrations of *CaCl*_2_. (d) Minimum distances between ACE2 and CoV-2 RBD as function of the residue index of ACE2 averaged over the trajectories from simulations at different concentrations of *NaCl*. (e) Minimum distances between ACE2 and CoV-2 RBD as function of the residue index of CoV-2 RBD averaged over the trajectories from simulations at different concentrations of *CaCl*_2_. (f) Minimum distances between ACE2 and CoV-2 RBD as function of the residue index of CoV-2 RBD averaged over the trajectories from simulations at different concentrations of *NaCl*. residue index

We then analyzed the root mean square fluctuation of the CoV-2 RBD - ACE2 complex at different salt concentrations of *CaCl*_2_ and *NaCl* (see Figure 3). As a general observation, we find that the N-terminal region between *α*1 and *β*4 of CoV-2 RBD fluctuates to the strongest extend with *RMSF* ≈ 1 nm. As another pattern, which we find in every RMSF colored structure, the binding interface between CoV-2 RBD and ACE2 fluctuates to a very low extent with *RMSF <* 0.3 nm. Another common feature is the high fluctation of the second helix of the N-terminal helical pair of ACE2 (helix pair”). Taking into consideration that there are high fluctuation patterns at low concentrations, which we find for *c*(*CaCl*_2_) = 0.06 M and *c*(*NaCl*) = 0.06 M (decreasing again for *c*(*CaCl*_2_) = 0.08 M and *c*(*NaCl*) = 0.08 M (see Figure 3 c, g)), we see a tendency for rising fluctuations with an increasing salt concentration, as we see in the RMSF colored structres for the concentrations *c*(*CaCl*_2_) = 0.3 M and *c*(*NaCl*) = 0.3 M (see Figure 3 b, f and e, i).

**Figure 3:**
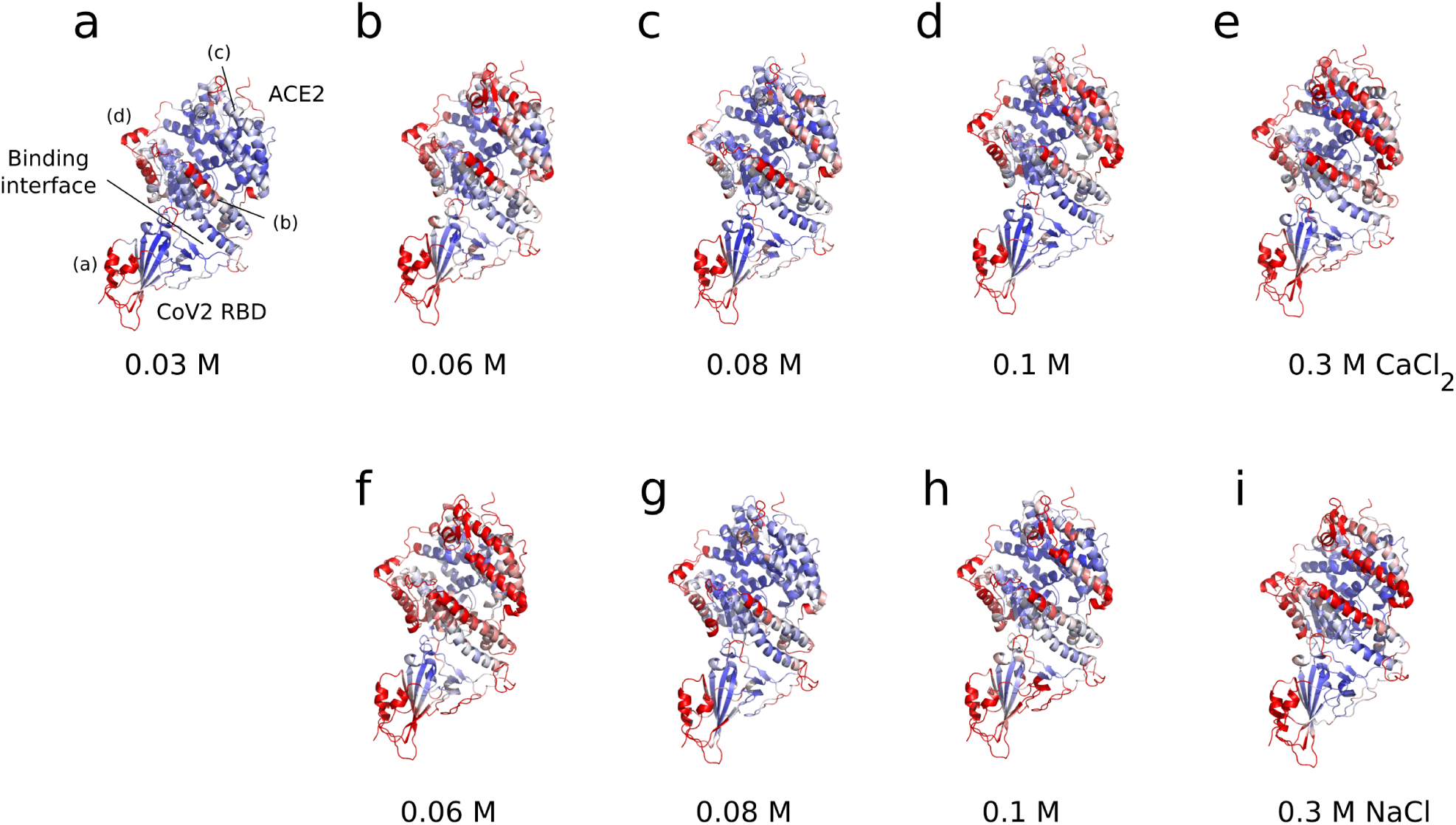
Color assigned root mean square fluctuation (blue : 0 nm, red : 1 nm) along the structure of the CoV-2 RBD - ACE2 complex in the *CaCl*2-systems (a-e) and in NaCl (f-i). (Subpanels : ((a)-(d)) in panel (a) : (a) N-terminal region of CoV-2 RBD. (b) N-terminal helix of ACE2. (c) Helical region”. (d) C-terminal lobe.)

### Interactions at the CoV-2 RBD - ACE2 interface at different salt concentrations

We analyzed the interaction energies at the CoV-2 RBD - ACE2 interface at two different concentrations of *CaCl*_2_ *c*(*CaCl*_2_) = 0.03 M) and *c*(*CaCl*_2_) = 0.3 M) and *NaCl c*(*NaCl*) = 0.06 M and *c*(*NaCl* = 0.3 M) as function of the residue index of ACE2 and CoV-2 RBD (see Figure 4). For the binding between CoV-2 RBD to ACE2, we observe larger fluctuations mainly due to the exchange of binding residues of CoV-2 RBD at ACE2. Interaction energies at the N-terminal region of ACE2 (i.e. the N-terminal helix), the relative log likelihood for binding ranges from -14.3 *k* _*B*_*T* to -0.86 *k* _*B*_*T* in the case of *CaCl*_2_ (see Figure 4 a). In the second binding region for residues in the range from # 320 to # 400 the interaction log likelihoods range from -13 *k* _*B*_*T* to zero representing the reference point. For the simulations at different concentrations of *NaCl* the energies are lower by approximately 1 *k* _*B*_*T* in the N-terminal region (see Figure 4 a and b). For both simulations, we find in general that the interaction log likelihoods decrease for higher salt concentrations by an approximate value ranging from 0.8 to 1 *k* _*B*_*T*. As a remark, we conclude here that the interchange of contacts at the interface between CoV-2 RBD and ACE2 increases at ACE2 with high salt concentrations, leading to a decrease in the interaction energy.

**Figure 4:**
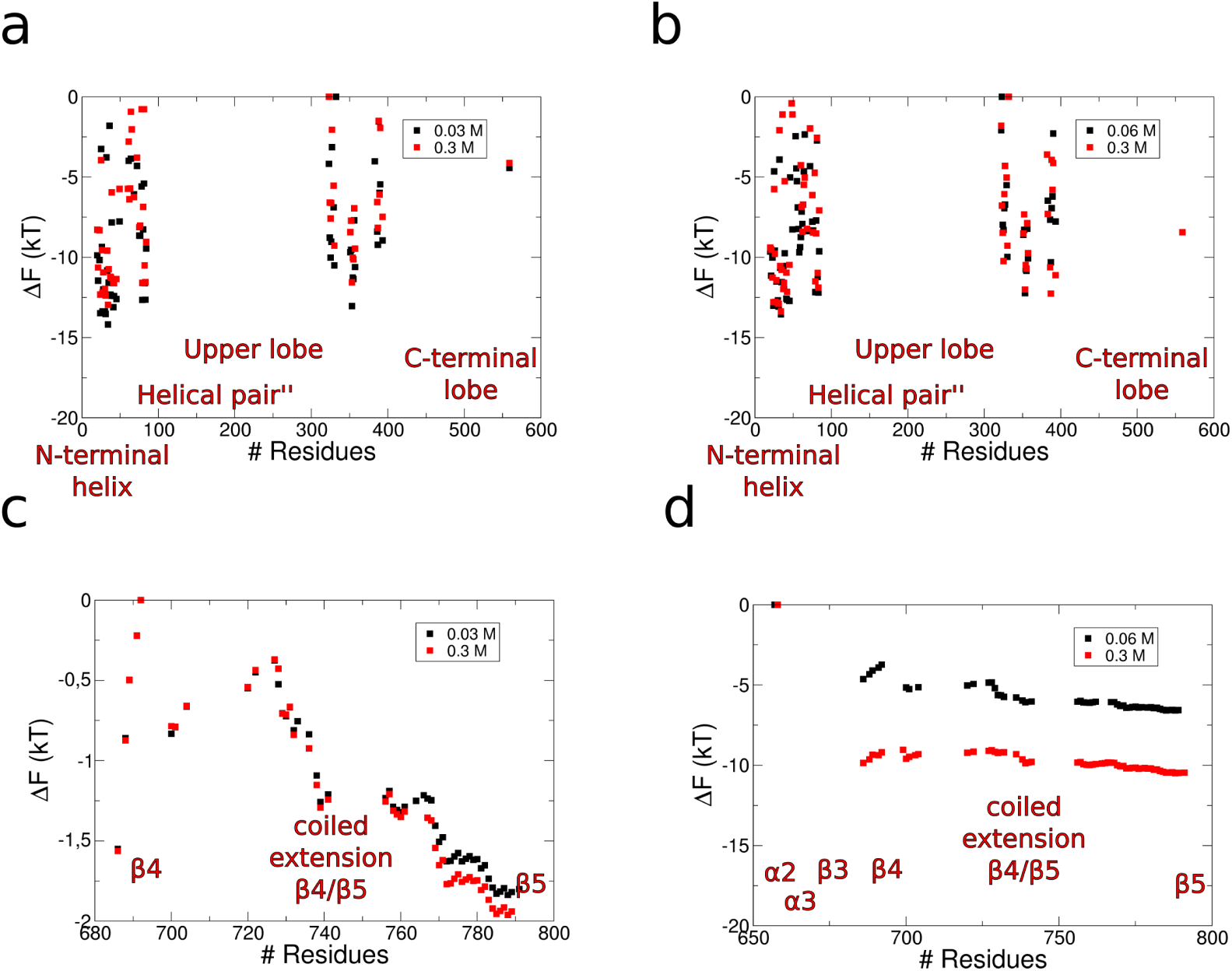
(a) Relative log likelihood of the interaction between CoV-2 RBD and ACE2 as function of the residue index of ACE2 at the concentrations 0.03 and 0.3 M *CaCl*2. (b) Relative log likelihood of the interaction between CoV-2 RBD and ACE2 as function of the residue index of ACE2 at the concentrations 0.03 and 0.3 M *NaCl*. (c) Relative log likelihood of the interaction between CoV-2 RBD and ACE2 as function of the residue index of CoV-2 RBD at the concentrations 0.03 and 0.3 M *CaCl*2. (d) Relative log likelihood of the interaction between CoV-2 RBD and ACE2 as function of the residue index of CoV-2 RBD at the concentrations 0.03 and 0.3 M *NaCl*.

In the opposite case, where we analyzed the relative log likelihood for binding of the CoV-2 RBD - ACE2 complex as function of the CoV-2 RBD index, we observe an increase of the affinity of CoV-2 RBD for ACE2 at an increasing electrolyte concentration (see Figure 4 c, d). For the simulations in *CaCl*_2_, we observe that the energy profile shows increasing binding energies at the coiled extension *β*4 / *β*5, while the energies for the higher concentration (*c CaCl*_2_) = 0.3 M) are shifted by 0.2 *k* _*B*_*T* towards higher energies (see Figure 4 c). For the simulation with *NaCl*, we find drastically higher interaction energies between -3.6 to -6.7 *k* _*B*_*T* at *c*(*NaCl*) = 0.06 M and energies between -8.9 to -10.4 *k* _*B*_*T* at *c*(*NaCl*) = 0.3 M, where we see an increase in the relative log likelihood for binding by 3 *k* _*B*_*T* with a salt concentration *c*(*NaCl*) = 0.3 M (see Figure 4 d).

### Conformations and dynamics of CoV-2 RBD - ACE2 at different salt concentrations

We compared the conformational changes depending from the salt concentrations using distance dependent maps averaged over the last frame of 50 ns in the simulations with *c(CaCl*_2_) = 0.03 and 0.3 M, and *c*(*NaCl*) = 0.06 and 0.3 M (see Figure 5). In each of the distance maps, we observe that the N-terminal helix of ACE2 binds to the coiled extension *β*4 / *β*5 of CoV-2 RBD. We also see that both domains are affected by internal conformational changes in each of the cases, while the largest internal reorientations can be found at *c* (*CaCl*_2_) = 0.3 M and *c*(*NaCl*) = 0.3 (see Figure 5 b, d). We observe strong reorientations in the regions of the Upper and the C-terminal lobe of ACE2, which are induced by the salt concentration. In terms of the relative values of the distance matrices, higher salt concentrations lead to a compaction between the helical pairs, the upper and the C-terminal lobe of ACE2, while the opposite occurs for the CoV-2 RBD part. In this case, we observe a decrease in the relative values along the contact pattern at high salt concentrations (see Figure 5 a-d). Although the internal dynamics of both domains change significantly, the binding between CoV-2 RBD and ACE2 is affected to a minor extent. Therefore, we performed a principal compontent analysis of each simulation (see Figure 6). The covariance analyses show similar patterns. Therefore, we only show the covariance matrix of the simulation with *c* (*CaCl*_2_) = 0.03 M (see Figure 6 a). We find specific patterns with higher principal deviations from the average structure at the Helical pairs, the central upper lobe of ACE2 and the N-terminal region (*α*1 - *β*4) of CoV-2 RBD (see Figure 6 a). When we analyzed the filtered trajectories using the first 8 eigenvectors of the covariance matrix, we find a general twisting mode of the N-terminal region of CoV-2 RBD and the helical pair” at the upper lobe of ACE2 (see Figure 6 b). Another principal motion of the N-terminal helical pair’ leads to a contraction with the helical pair”, while parts of the C-terminal lobe fluctuate perpendicular into the bulk (see Figure 6 b). In contrast to these larger modes, the binding interface between CoV-2 RBD and ACE2 remains unaffected.

**Figure 5:**
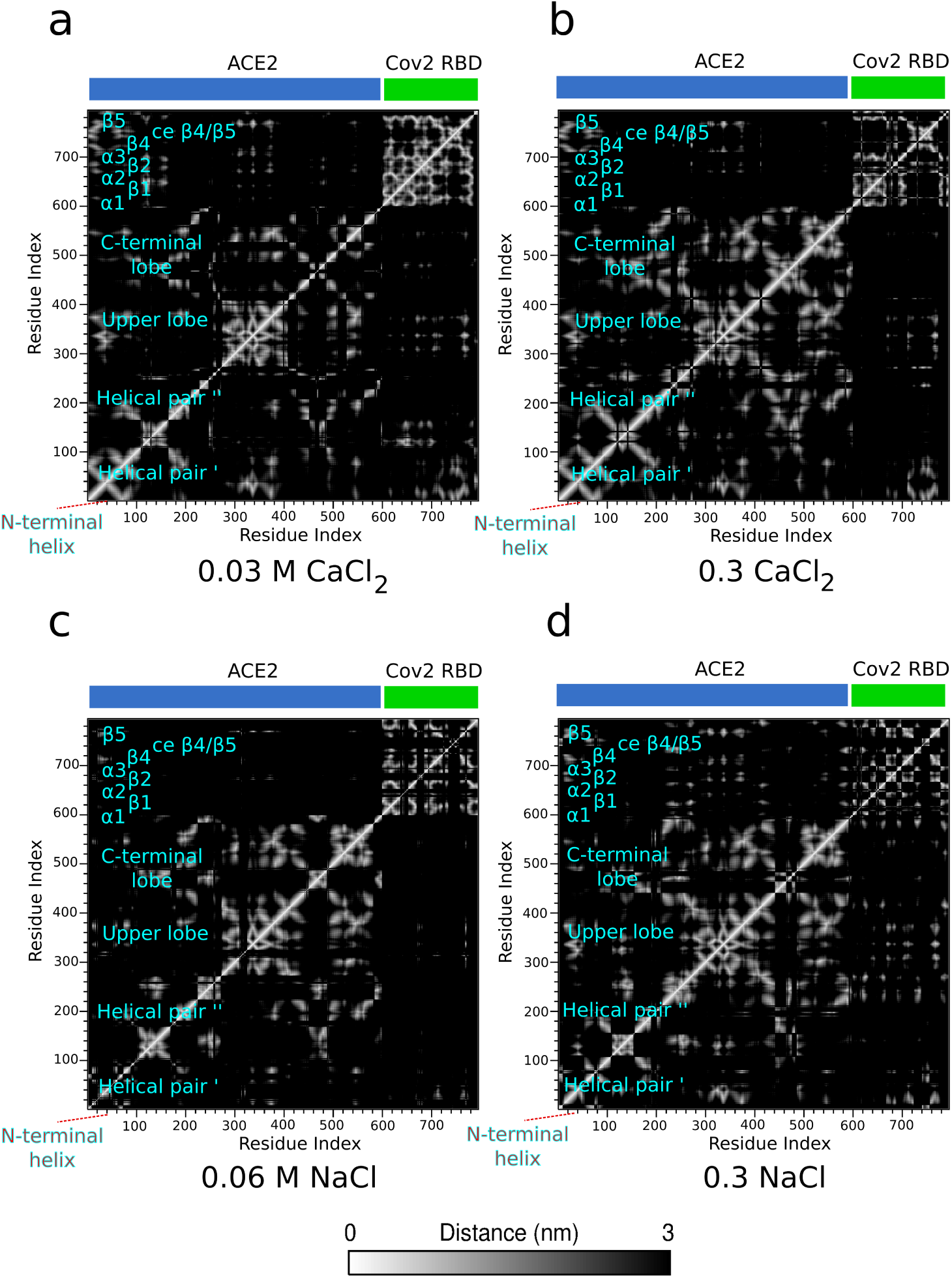
Distance maps averaged over the last 50 ns of the 750 ns trajectories from simulations with (a) 0.03 M *CaCl*_2_, (b) 0.3 M *CaCl*_2_, (c) 0.06 M *NaCl* and (d) 0.3 M *NaCl*.

**Figure 6:**
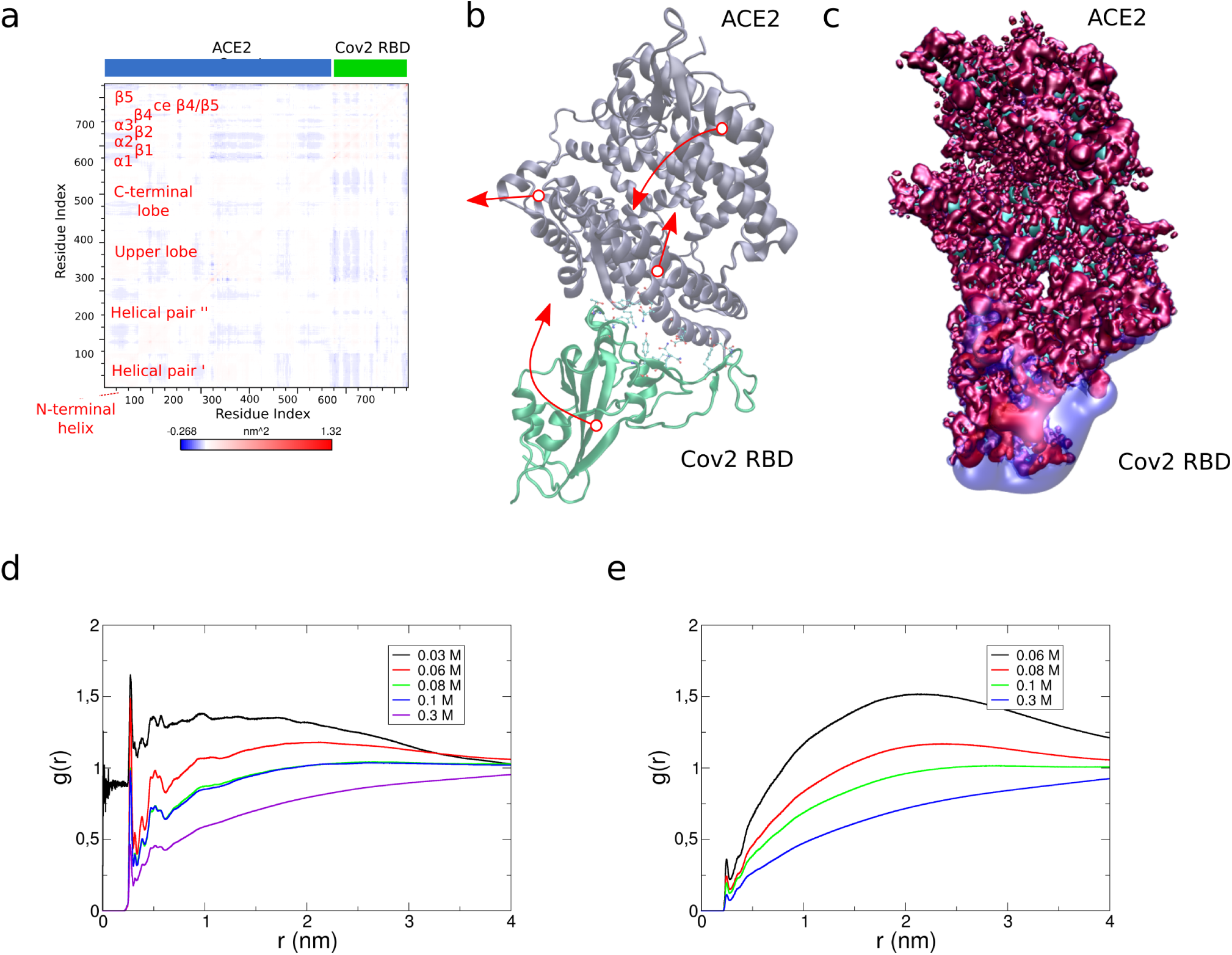
(a) Covariance matrix from the simulation at 0.03 M *CaCl*_2_. (b) Principal motions of the CoV-2 RBD - ACE2 complex extracted from the covariance analysis. (c) Electrostatic energy surfaces at +1 (red) and -1 (blue *k* _*B*_*T / e* from APBS calculations of the CoV-2 RBD - ACE2 complex. (e) Radial distribution functions of the total number of *CaCl*_2_ electrolytes in the system in relation to the protein complex. (f) Radial distribution functions of the total number of *NaCl* electrolytes in the system in relation to the protein complex.

Finally, we analyzed the electrostatic patterns of the CoV-2 RBD - ACE2 complex and the radial distribution functions between the protein complex and the electrolytes (see Figure 6 c). To our surprise, we found that the electrostatic potential of ACE2 at the helical pairs, the upper and the C-terminal lobe are almost charge neutral considering the two plotted electrostatic potentials (±1 *k* _*B*_*T / e*). In contrast to that, we see a stronger negative polarization in the region of *α*1, *β*1, *α*2 and *α*3 at the N-terminal region of CoV-2 RBD as well as along the binding interface at the coiled extension *β*4 / *β*5 (see Figure 6 c). In the radial distribution functions, we do find a systematic decrease of the probability density near the surface of the protein complex starting at low concentrations in the range from *c* = 0.03-0.08 M (see Figure 6 d, e). That can be interpreted as a decrease in the total probability density to localize electrolytes at the surface of the protein in relation to the total number of ions in the system. In that case, we applied no normalization of the RDF, which leads to values bigger than 1 for radii in the bulk. We interpret this effect as a saturation effect of ions adsorbed to the surface of the protein already at low concentrations, while a larger fraction of ions resides in the bulk and only imposes a fluctuating electric field on the protein complex.

## Discussion

As expressed first in the famous Hofmeister series, electrolytes affect biomolecular processes through specific and non-specific interactions with the biomolecule (43–45). Franz Hofmeister discovered that the effectiveness of salts for protein precipitation generally followed a specific order, regardless of the protein being investigated (46). The specific Hofmeister series of effects of electrolytes on proteins has been extended to effects on the surface tension, the surface potential (47, 48), and salt effects on a variety of macromolecular processes, such as micelle formation, “salting out“-behavior, the effect on nonpolar compounds, and protein folding (49–52). A number of ions as part of the Hofmeister series favour protein folding (53), while specific electrolytes can lead to denaturing conditions. The theories on how electrolytes affect the stability of proteins are explained by two different types of interaction : The interaction of ions with polar and non-polar groups of a protein. For example, the salting-out effect on proteins containing nonpolar groups has been explained by a cavity model (44). That model defines surface tension increments upon binding of an ion to a protein and predicts observations of increasing salting-out constants, which is proportional with the number of hydrophobic carbon atoms in the aliphatic side chains of a protein. However, the mechanism of the interaction between ions and a protein is still not well understood, and it is controversial whether the ion-protein interactions are ion-specific. The interactions can be not ion-specific, if no polar groups are present and a potential specificity resides in interactions with nearby nonpolar groups. For example, a non-specific salting-in interaction can occur between monovalent ions and dipolar molecules, where the energy depends from the ionic strength and is independent from the position of an ion in the Hofmeister series. In that case the strength of the interaction is dictated by the ionic strength and the role of water remains one of the key factors in the understanding of changes in the stability of proteins. Although a broad knowledge exists on the changes of structure and dynamics of water through the addition of electrolytes (54, 55), an isolated view on ion-protein interactions in the Hofmeister series is broadly accepted by the community, while protein hydration seems to be studied separately and independent from the effect of electrolytes (56).

Generally speaking, we do not find distinct changes in the behavior of the CoV2 RBD - ACE2 complex with changing salt concentrations, such as a cleavage of the complex or partial unfolding within the simulated time. We state that higher ionic concentrations beyond physiological salt concentrations can shift the equilibria towards disassembly of the protein complex, or stabilize the interaction (57). As a general remark, we observe that the salt affects the internal dynamics of CoV2 RBD and ACE2, where we see increasing fluctuations and a compaction, while a significant stabilizing effect on the interaction between CoV2 RBD and ACE2 with rising salt concentration can only be observed with sodium chloride. For high concentrations of *c*(*CaCl*_2_) = 0.3 M, the increase in the interaction energies remains below 0.5 *k* _*B*_*T*. We emphasize that this stabilization effect occurs with no major conformational change of the entire complex within the simulation time.

ACE2 is a protein bound to the membrane of human cells, where comparatively strong electrolyte effects play an imminent role (58). Physiological salt concentrations near the membrane are at an approximately molar concentration of 0.5 M, where the intracellular level can be 100-1.000 times lower than in the extracellular environment (59). Thus, the formation of the protein complex between CoV-2 RBD and ACE2 occurs within a highly polar environment. In our simulations at varying salt concentrations of two different electrolytes, we observe that the contact between both proteins remains stable, although we observe increasing internal structural rearrangements, such as a compaction of ACE2 at higher salt concentrations. We also find structural changes for CoV-2 RBD, while the binding site between CoV-2 RBD and ACE2 remains stabilized and compact. We find that major fluctuations occur at the N- and C-termini of CoV-2 RBD and at the peripheral helical pairs of ACE2. However, the bending motion of regions in ACE2 and the terminal sections of CoV-2 RBD conserve the interface between both proteins. While we find that ACE2 contains an neutral electrostatic potential, CoV-2 RBD contains a strongly negative electrostatic potential lobe. Although we see that the adsorption patterns of salt change at the surface of the protein with varying salt concentrations, the interaction of the electrolytes at the surface leads to higher fluctuations of the protein complex leading to a slight increase of the relative log likelihood of binding between CoV-2 RBD and ACE2. We observe that the regions at the N- and C-termini of CoV-2 RBD and the two helical pairs of ACE2 are governed by increasing fluctuations, which, however, do not perturb the binding interface. We speculate that one of the reasons for this stabilization is the evolutionary adaption of CoV-2 to homologous ACE2 proteins in other mammal organisms (19, 60).

## CONCLUSIONS

In this article, we investigated the effect of electrolytes on the stability of the complex between the coronavirus 2 spike protein receptor domain (CoV-2 RBD) and ACE2. The formation of this protein complex plays an important role in the activation cascade at the viral entry of CoV-2 into human cells. Near the cellular surface, where the activation of CoV-2 occurs, electrolyte conditions play an important role, especially in the interaction of proteins. As another aspect, the binding interface of the CoV-2 RBD - ACE2 complex is highly hydrophilic. We performed long-time MD simulations of the CoV-2 RBD - ACE2 complex at varying salt concentrations over the concentration range from 0.03 M to 0.3 M of calcium and sodium chloride over an individual simulation length of 750 ns in 9 independent simulations (6.75 *µs* total). We observed that the CoV-2 RBD - ACE2 complex is stabilized independent of the salt concentration. We identified a strong negative electrostatic potential at the N-terminal part of CoV-2 RBD and we observed that CoV-2 RBD binds even stronger at higher salt concentrations. We found that the dynamics of the N-terminal part of CoV-2 RBD stabilize the protein complex leading to strong collective motions and a stable interface between CoV-2 RBD and ACE2. We speculate that the sequence of CoV-2 RBD might be optimized for a strong binding to ACE2 at varying salt concentrations at the cellular surface, such that CoV-2 activation is possible within a wide range of electrolyte concentrations.

